# Pre-saccadic neural enhancements in marmoset area MT

**DOI:** 10.1101/2022.10.11.511827

**Authors:** Shanna H. Coop, Jacob L. Yates, Jude F Mitchell

## Abstract

Each time we make an eye movement, attention moves before the eyes, resulting in a perceptual enhancement at the target. Recent psychophysical studies suggest that this pre-saccadic attention enhances the visual features at the saccade target, whereas covert attention causes only spatially-selective enhancements. While previous non-human primate studies have found that pre-saccadic attention does enhance neural responses spatially, no studies have tested if changes in neural tuning reflects an automatic feature enhancement. Here we examined pre-saccadic attention using a saccade foraging task developed for marmoset monkeys. We recorded from neurons in the middle temporal (MT) area with peripheral receptive fields that contained a motion stimulus which would either be the target of a saccade or a distracter as a saccade was made to another location. We established that marmosets, like macaques, show enhanced pre-saccadic neural responses for saccades towards the receptive field, including increases in firing rate and motion information. We then examined if the specific changes in neural tuning might support feature enhancements for the target. Neurons exhibited diverse changes in tuning, but predominantly showed additive and multiplicative increases that were uniformly applied across motion directions. These findings confirm that marmoset monkeys, like macaques, exhibit pre-saccadic neural enhancements during saccade foraging tasks with minimal training requirements. However, at the level of individual neurons, the lack of feature-tuned enhancements is similar to neural effects reported during covert spatial attention.

**Significance Statement:** Attention leads eye movements producing perceptual enhancements at saccade targets. Recent psychophysical studies indicate that increases in pre-saccadic sensitivity are concentrated around features of the target. We tested at the neural level how pre-saccadic attention modulates the tuning curves of visual neurons in area MT of marmoset monkeys. While neurons exhibited clear pre-saccadic enhancements that were consistent with previous studies in macaques, the changes in tuning were uniform across tuning. These results show pre-saccadic enhancements are a general feature of visual processing, shared by New World monkeys, but at the level of individual neuron’s enhancements are uniform across features much like what has been reported previously for covert attention.

## Introduction

Visual attention is strongly linked to eye movement planning and saccadic control (Bisley, 2011; Squire et al., 2013). Every saccade is preceded by a shift of attention that enhances perception of the saccade target, called pre-saccadic attention (Kowler et al., 1995; Deubel & Schneider, 1996; Rolfs et al., 2011; White et al., 2013). Pre-saccadic attention is automatic and occurs rapidly within 50-100ms before saccades (Deubel, 2008; Rolfs et al., 2011; Rolfs & Carrasco, 2012; Li et al., 2016; Ohl et al., 2017). It also appears to be obligatory occurring even when it is detrimental to task demands (Montagnini & Castet, 2007; Deubel, 2008; Steinmetz & Moore, 2010). Neural studies with non-human primates have shown that pre-saccadic attention involves enhancements in firing that increase neural sensitivity, and to a first approximation, are highly similar to that seen during covert attention where the eyes remain at central fixation (Steinmetz & Moore, 2010; Squire, et al., 2013). However, psychophysical studies highlight ways in which pre-saccadic attention differs from covert attention (Li et. al., 2021a,b). The differences between these mechanisms at the neural level remains uncertain.

Recent human psychophysics suggest that pre-saccadic attention involves a concentration of enhancement around the saccade target’s features (Li et al., 2016; Ohl et al., 2017; Li, et al., 2021a). The concentration of sensitivity around a target feature could be implemented at the neural level by feature gain, similar to a feature-based attention. Neurophysiology studies of feature-based attention have found that neurons sharing features with an attended feature increase the gain of their responses while those selective to opposite features are suppressed (Treue & Martinez-Trujilo,1999; Martinez-Trujilo & Treue, 2004). By contrast, a pure spatial selection of the target, as in covert spatial attention, is associated with an increases in the gain that is uniform across features (McAdams & Maunsell, 199a). If pre-saccadic attention engages automatic selection of target features, we would predict feature-specific gain rather than spatial gain. To test between these alternatives requires detailed measurement of neural tuning.

Previous studies of pre-saccadic attention in macaques have found improvements in neural sensitivity but without careful examination of feature tuning. It is known that visual neurons in several brain areas increase firing rates and stimulus selectivity during pre-saccadic attention when a saccade is planned to a stimulus within a neuron’s receptive field (Moore et al., 1998; Li & Basso, 2008; Moore & Chang 2009; Steinmetz & Moore 2010; Merrikhi et al, 2017). It has also been shown at off-target locations that there are enhancements when the RF stimulus matches features of the saccade target (Burrows et al, 2014). However, no studies have examined if changes in tuning curves might reflect feature or spatial gain.

We examined how pre-saccadic attention modulates tuning curves in areas MT/MTC of the marmoset monkey. The marmoset is a small-bodied New World primate that has gained interest for neural investigations due to feasibility of genetic manipulation (Belmonte et al., 2015), and advantages for imaging and array recordings (Solomon & Rosa, 2014; Mitchell & Leopold, 2015). Area MT/MTC neurons exhibits similar tuning for motion direction as macaques (Elston & Rosa, 1999). However, it remains untested if marmosets share similar mechanisms of visual attention. In the macaque it is established how MT neurons are modulated by feature-based attention (Treue & Martinez-Trujilo, 1999; Treue & Martinez-Trujillo, 2004), and also that they show pre-saccadic enhancements (Merrikhi et al, 2021). We developed a saccade foraging paradigm for marmoset monkeys to test for changes in neural tuning under pre-saccadic attention. We first established that, like macaques, marmoset neurons show pre-saccadic increases in neural firing and sensitivity for motion direction. Then by varying motion direction independently across trials we sampled full tuning curves for motion direction and tested if they exhibit feature-specific enhancements for the saccade target.

## Results

We used a saccade foraging task to measure the modulation of neural firing and motion tuning during pre-saccadic attention **(Figure 1A)**. In each trial, the monkey was trained to maintain fixation on a central point for 100-300 ms, after which three random dot field motion stimuli appeared in peripheral apertures of equal eccentricity and separation from each other. The monkey responded by making a saccade to one of the three apertures immediately after stimulus onset. While monkeys performed this foraging task, we recorded from individual neurons in visual areas MT/MTC **(Figure 1B)**. The apertures were positioned such that one of them was centered inside the receptive field of the neurons under study. Thus, across trials the monkeys performed saccades either towards the receptive field (“Towards” condition) or away from the receptive field (“Away” condition). Because we examined responses in the pre-saccadic epoch while the eyes were still at fixation, the sensory stimuli were matched between these conditions so we could isolate the effects of saccade planning on neural responses. Our goal was to measure how neural tuning curves differ for saccades towards the receptive field as opposed to away from it.

**Figure 1.**
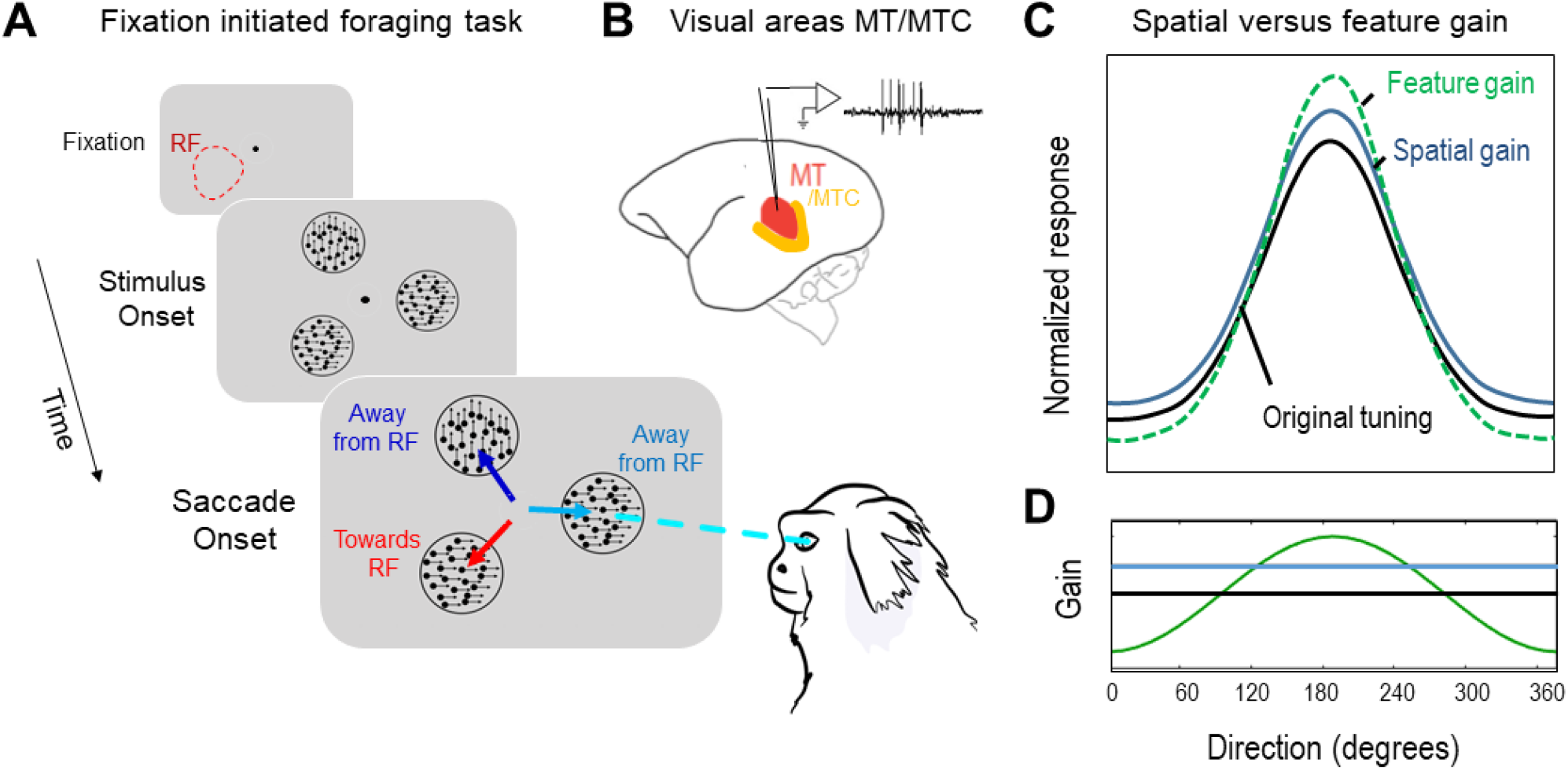
A saccade foraging task to test if pre-saccadic attention involves feature-gain. **(A)** The saccade foraging task encouraged sampling between three peripheral apertures, one at the RF location. Subjects fixated centrally for 100-300 ms after which dot-motion apertures appeared, and reward was given for making a saccade to an aperture that differed from the previous trial. Apertures contained 100% coherent moving dots and across apertures the motion direction was sampled independently. **(B)** Neural recordings were made from areas MT/MTC to measure neural tuning curves for motion direction. **(C)** Example tuning curves demonstrate changes predicted for a spatial gain model (blue) and the feature-similarity gain model (dashed line, green), compared against the neutral tuning curve (black). **(D)** The proportional gain as a function of direction would be predicted to be uniform for the spatial gain model (blue). For feature-gain it should have a peak at the preferred direction relative to the flanks (green).

Recent studies in human psychophysics suggest that pre-saccadic attention differs from covert attention in important ways, specifically involving an automatic narrowing of sensitivity around the feature of the saccade target, whereas covert spatial attention applies a uniform gain at the attended location independent of tuning. A narrowing in feature sensitivity could manifest at the level of individual neuronal tuning curves in a variety of ways. For example, in covert attention tasks previous studies have identified individual MT neurons that change their tuning according to widely used gain models, including spatial gain (McAdams & Maunsell, 1999a) and feature-similarity gain as observed when feature-based attention is involved (Treue & Martinez-Trujilo, 1999; Treue & Martinez-Trujillo, 2004). If presaccadic attention only enhances spatial gain there will be a uniform multiplicative increase across all motion directions **(blue curve, Figure 1C)**. Considering how that would impact the proportional gain across the tuning curve such a spatial gain would give constant positive value (flat line) across directions **(Figure 1D)**. Alternatively, the feature-similarity model would preferentially increase the gain for preferred motion directions while suppressing gain for non-preferred directions **(green curve, Figure 1C)**. In that case the proportional gain would not be uniform across the curve but rather show an enhancement at the peak of tuning and suppression away from the peak **(green curve, Figure 1D)**. We designed our saccade foraging task to sample across trials different motion directions in the pre-saccadic epoch of response to reconstruct full neural tuning curves and test if changes in tuning favor feature specific gain as opposed to spatial gain.

### Measurement of MT/MTC receptive fields during free viewing

To properly position peripheral stimuli in the saccade foraging task we must first determine the receptive fields of MT/MTC neurons. As marmosets are less able to maintain fixation on central locations for extended periods during receptive field mapping (Mitchell et al., 2014; Yates et al., 2021), this presented a unique challenge. We developed a novel free-viewing approach to map the receptive fields of MT/MTC neurons (Yates et al., 2021; see methods). In brief, marmosets were allowed to explore a full-field display of moving dots, with 4-16 dots displayed at any time and each dot being 1-degree visual angle (dva) in diameter. The dots flashed at random locations and then moved along a single direction of motion at 15 dva/sec for a duration of 50 ms **(Figure 2A)**. Off-line we corrected for eye movements during viewing of the full-field stimulus and reconstructed the stimulus history in a retino-topic coordinate frame to assess visual receptive fields. Those visual locations that exhibited responses significantly above the pre-stimulus baseline time-locked to a dot appearing were labeled to identify the receptive field **(Figure 2B, top)**. Then the responses to dots within the receptive field were further broken down based on their direction of motion to estimate the neuron’s motion tuning **(Figure 2B bottom)**. The example cell illustrated had a visual latency of 40-50 ms with a receptive field in the lower left visual quadrant and strong motion tuning as reflected by a Direction Selective Index (D.S.I.) significantly above zero, consistent with a typical MT/MTC neuron.

**Figure 2.**
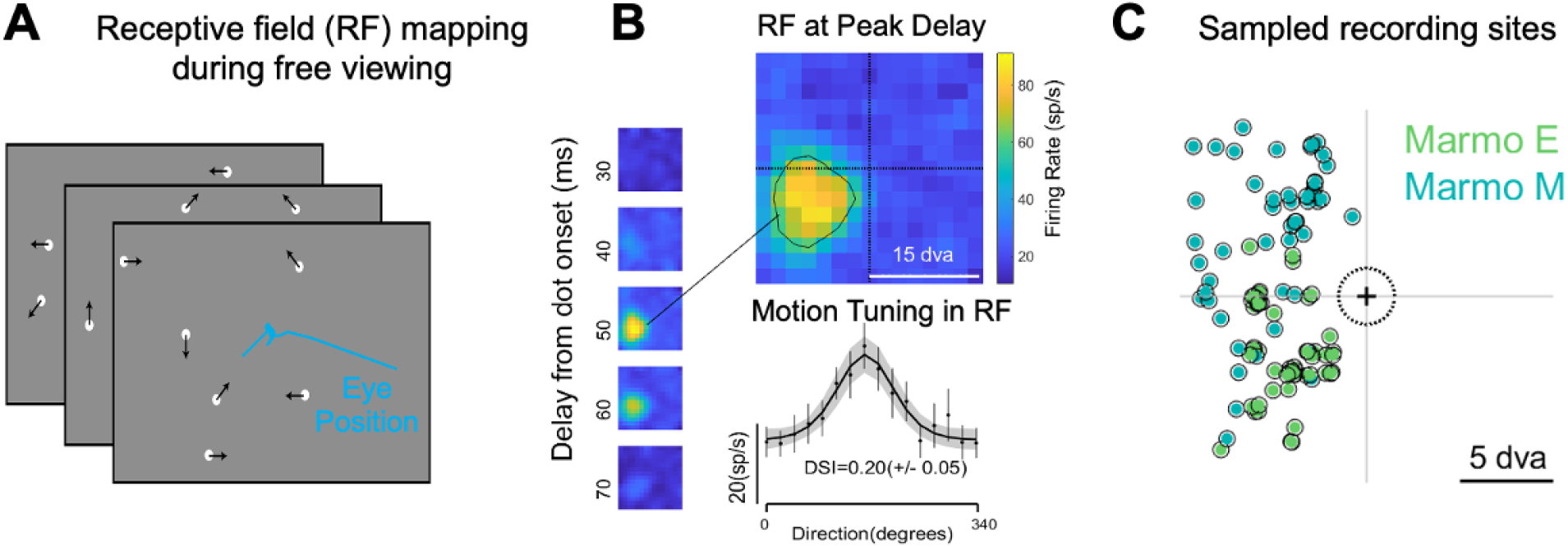
Receptive field estimation during free-viewing in marmoset MT/MTC. **(A)** Marmosets freely view a full-field sparse motion-noise stimulus consisting of flashed dots that onset at random locations and move for 50 ms before replotting. **(B)** Off-line correction for eye movements enabled us to reconstruct the spatial/temporal receptive field and motion tuning curve for a motion selective neuron. Error bars in the tuning curve are 2 S.E.M. **(C)** The distribution of spatial receptive fields sampled across recordings in both animals: Marmoset E (green) and Marmoset M (blue).

We recorded single and multi-unit activity from two marmoset monkeys across 38 and 52 experimental sessions respectively. Neural recordings were targeted for Middle Temporal (MT) area based on retinotopy and motion selectivity, but some neurons with receptive fields near the vertical meridian may have been included from adjacent Middle Temporal Crescent (MTC) area, which like area MT has a significant portion of neurons with motion selective responses and comparably sized receptive fields (Rosa & Elston, 1998). We only included neurons for analyses if they exhibited a visual response with significant motion tuning and a minimum evoked firing rate of 1 spike/sec (see methods). The first marmoset monkey (Marmoset E) was recorded using single tungsten electrodes during initial studies while advanced array recordings methods were still in development. We obtained 116 units of which 87% showed significant visual responses, and of those 75% had significant motion tuning, giving 73 units in total (39 single units, 34 multi-units). The second marmoset monkey (Marmoset M) was recorded after we had refined our recording methods to include a 64-channel linear array yielding higher cell counts. We obtained 872 units of which 60% had significant visual responses, and of those 90% which had significant motion tuning, giving 472 units in total (444 single units, 28 multi-units). In all subsequent analyses the two marmosets are presented separately, first as the second animal would dominate the sample based on neuron numbers, but also because the different recording methods impose different sampling biases across neuron types and cortical layers.

The distribution of visual field locations sampled for individual recording sessions across the two marmosets covered the upper and lower left hemi-field **(Figure 2C)**. For array recordings, we typically isolated several neurons in a single recording but because the marmoset cortex is smooth our linear arrays were oriented down a single cortical column, and thus had largely overlapping receptive fields. We thus were able to test a single visual field location that encompassed different neurons on the same array. We sampled from the left hemi-field in both animals. Biases towards lower or upper visual fields varied between animals because of the position of blood vessels in their tissue, which we avoided in placing electrodes. Despite variation in electrode placement and recording methods, we find similar qualitative patterns of neural modulation across the two animals.

### Marmoset behavior in a saccade foraging task for study of pre-saccadic attention

While the smooth cortical surface of marmosets facilitated access of areas MT/MTC for neural investigation, a key disadvantage of working with marmosets is the number of trials they can perform in highly constrained behavioral tasks (Mitchell et al., 2014). In the macaque, studies of pre-saccadic attention have imposed constraints to maintain central fixation that are similar to covert attention task, with an extended period of fixation prior to making a saccade to a cued target (Moore & Chang, 2009; Steinmetz & Moore, 2014). Here we took a different approach with marmoset monkeys to instead optimize the number of saccade trials while minimizing the duration of fixation, and thus the total duration of individual trials. Marmosets completed 413 and 578 trials on an average session that met criteria for obtaining accurate initial fixation for a brief fixation epoch (1.5 dva fixation window) and then making an accurate saccade targeting one peripheral aperture (see methods). To encourage foraging between different locations across trials, a juice reward was delivered if the animal selected an aperture that differed from that selected in the previous trial. Initial piloting of the task demonstrated that marmosets foraged for more trials when the task included three apertures instead of two. During each recording session we positioned one of the three apertures over the neural receptive fields under study. By encouraging foraging between locations, we were able to sample neural responses both when saccades were made towards or away from the receptive field.

Marmoset monkeys acquired fixation accurately to initiate trials and sample across apertures locations during the foraging. The fixation and saccade end-points from a typical behavioral session are illustrated in **Figure 3A**. The color indicates which aperture location was selected in each trial, with red indicating saccades towards the RF and dark or light blue locations away from the RF. Zooming in on the period of central fixation, the 2D eye position clustered within the fixation window during the 100 ms epoch preceding saccade onset and overlapped regardless of the target selected **(Figure 3B)**. The overlap in central fixation was consistent across sessions and the two monkeys when comparing between conditions where the saccade was towards (red) or away (blue) from the receptive field **(Figure 3C)**. Across trials monkeys sampled across all three aperture locations as illustrated by the distribution of saccade end-points for the example session **(Figure 3A)**. Average trial counts favored sampling of the RF location (red) in both animals reflecting an unintended alignment in their spatial biases **(Figure 3D)**. Although marmosets foraged different locations across trials, they were not perfect at avoiding a return to the location selected in the previous trial **(Figure 3D, filled regions)**. Their return saccades to previous locations reflected the sampling bias towards the RF location. The biases towards the RF were further reflected in the distribution of saccadic latencies. Both monkeys made faster saccades towards the RF location (shown in red) as compared to the away locations (shown in blue) with median latency shifts of 10 and 15 ms respectively **(Figure 3E)**. To control for differences in the saccadic timing between towards and away conditions, all subsequent analyses comparing the firing rates between those conditions first match the saccadic latency distributions by resampling the trials from each session (see methods). This enables us to match saccade timing in addition to sensory conditions between towards and away conditions for further analyses.

**Figure 3.**
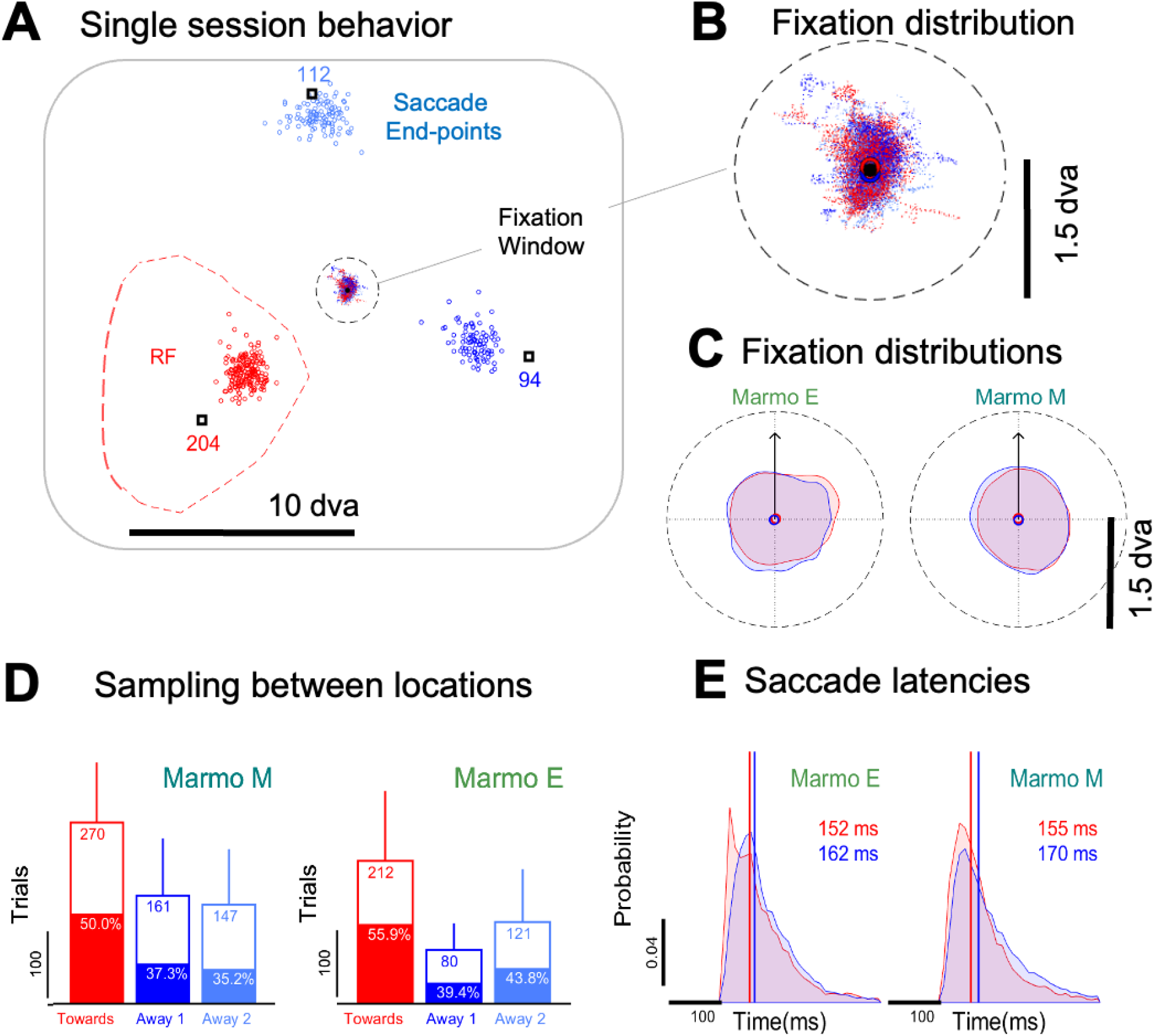
A saccade foraging task enables sampling between targeted locations. (**A)** The 2D eye position clustered around fixation before saccade onset. The color at fixation indicates which target location was subsequently selected (red, towards RF; blue and light blue, away from RF). Black square represents the aperture centers, with the saccade end-points shown in color. The receptive field of the neuron recorded overlapped one of the targets (red dotted line). (**B)** A close-up of the 2D eye position distribution during fixation for the example session. **(C)** Fixation distributions overlapped for the Towards RF (red) and Away RF conditions (blue) after being aligned to the saccade direction (saccade direction indicated by the arrow, contours indicate the 95% circumscribing region). **(D)** The histogram of trials across sessions reflects that all locations were sampled, with filled regions indicating the proportion of trials where saccades returned to the same location in consecutive trials (error bars one standard deviation). **(E)** The distribution of saccade latencies for each animal averaged across sessions for the Towards RF (red) and Away RF conditions (blue).

### Marmoset monkeys exhibit single unit neural signatures of pre-saccadic attention

As illustrated for a single example cell, the firing rate in pre-saccadic epoch increased for saccades made towards the RF location **(Figure 4)**. Spiking was time-locked across trials to the saccade onset for the Towards RF condition (in red) and Away RF conditions (in blue) **(Figure 4A)**. Averaging firing across the trials the mean rate showed no significant rate modulation when time locked to stimulus onset but had grown towards a modest increase around the time of saccade onset **(Figure 4B)**. The response at the stimulus onset from 40-100 ms did not exhibit a significant difference for Towards vs. Away conditions (58.5 vs 59.8 sp/s; Ranksum, p = 0.7625). The rate had a modest increase approaching significance for the Towards condition from −30 to 30 ms around the saccade onset (49.9 vs. 41.5 sp/s; Ranksum, p = 0.056). The mean firing rate, however, is averaged across trials that included different stimulus motion directions that sampled from 16 different directions from 0 to 360 degrees. When instead breaking out changes in rate as a function of direction **(Figure 4C)**, we observed a highly significant increase in the gain of the tuning curve of the example neuron in the Towards as compared to Away condition (Towards Amp: 100.4, Away Amp: 76.26; Z-transform based on von Mises fit confidence intervals, p = 0.00024).

**Figure 4.**
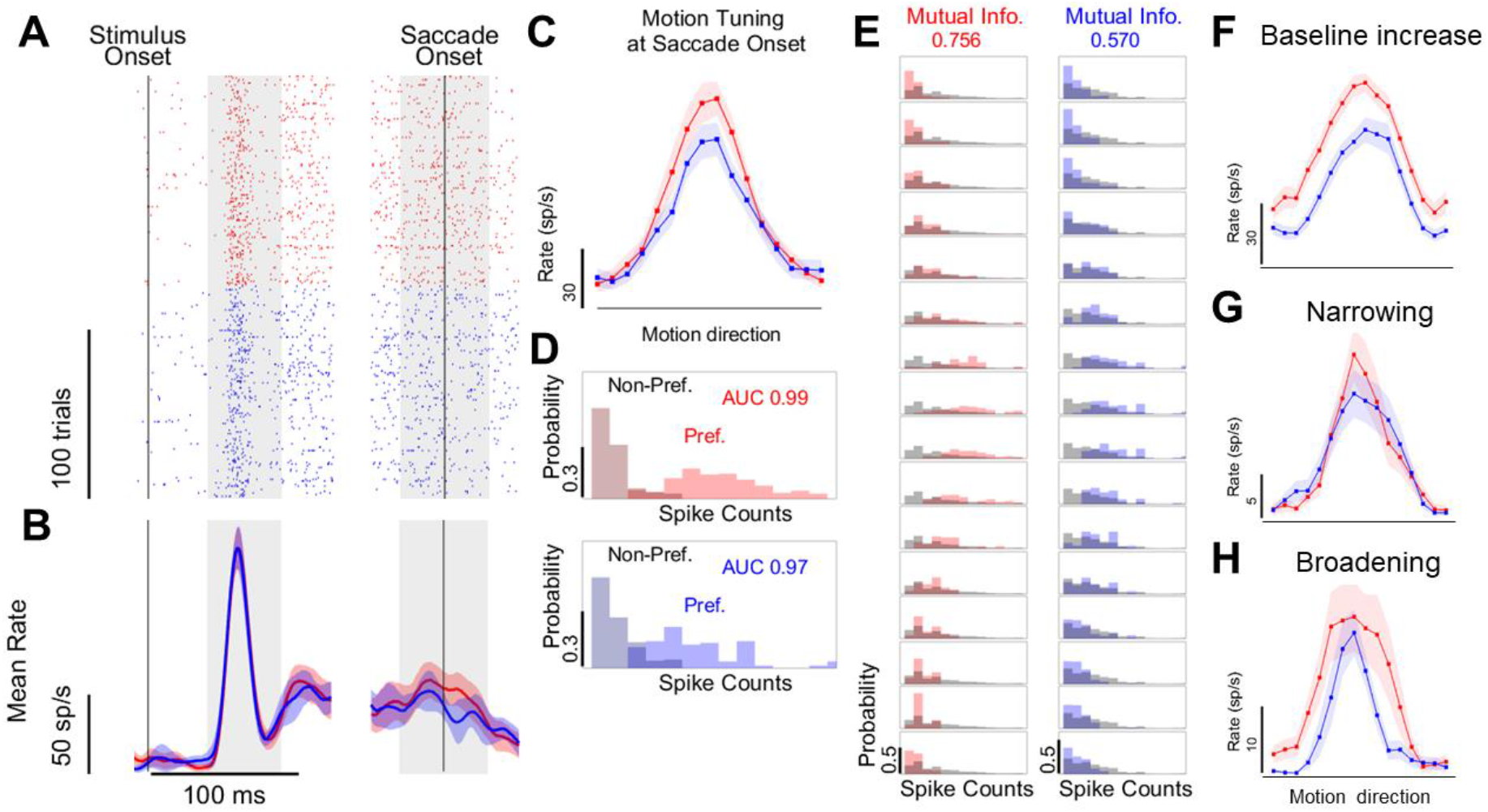
Example MT/MTC cells show a variety of enhanced pre-saccadic firing. **(A)** On the left, spike rasters are aligned to either stimulus onset or saccade onset for “Towards RF” (red) and “Away RF” trials (blue). Trials for each condition are arranged in ascending order by saccade latency. **(B)** The mean firing rate averaged across trials for each condition is shown. Error bars reflect 2 S.E.M. **(C)** The mean firing rate shown as a function of motion direction during the saccade onset interval (−30 to 30 ms around the saccade onset) reflects a gain increase for the Towards (in red) condition (error bars are 2 S.E.M). **(D)** The distributions of spike counts for the non-preferred and preferred motion directions in the two conditions. Neural sensitivity can be assessed by the area under the curve (AUC) of the receiver operating characteristic curves (ROC), but for this neuron does not differ substantially despite the large increase in gain. **(E)** The mutual information measure of sensitivity uses the spike count distributions per motion direction (shown in color) compared to the marginal distribution (shown in grey) and reflects an increase for the example neuron. **(F-H)** Tuning curves (same conventions as C) illustrating other neural changes in tuning.

Previous studies have reported increases in neural sensitivity during pre-saccadic attention based on comparing responses to preferred and non-preferred stimuli (Moore & Chang, 2009; Steinmetz & Moore, 2010). To assess the sensitivity of neurons to stimulus motion based on their tuning curve we instead computed the mutual information (M.I.). The mutual information provides a measure that is similar to the area under the curve (AUC) in the receiver-operate characteristic (ROC) function, but generalized for a complete tuning curve rather than just the case of two stimuli (Hastopolous et al., 1998). Similar to AUC, the mutual information depends not only on the separation in firing rates across the different stimulus conditions, but also the variability in responses, which describes neural sensitivity based on how well the spike count distributions for different stimuli can be separated. For the example neuron there was almost no difference in the AUC measure for the Towards (0.99) versus Away (0.97) condition **(Figure 4D)**. This simply reflects a saturation in that measure as the preferred and non-preferred stimuli in the tuning curve were extremely well separate in response. However, the mutual information remains sensitive to the separation across the entire curve and reflects an increase from 0.570 to 0.756 bits of information **(Figure 4E),** which approached significance for this neuron (Z-transform based on confidence intervals, p =0.065).

Across the population we found a diverse patterns of modulation in tuning curves. Similar to what has been observed in covert attention, we find that some cells show an increase in baseline firing rate with attention **(Figure 4F)**, while others show an increase in gain **(Figure 4C)**. However, we also find example cells with a narrowing in width **(Figure 4G)** that would be consistent with a feature-based gain. However, there are also many neurons that exhibit the opposite pattern, showing a broadening in half-width **(Figure 4H)**. Thus, we sought to determine how the population changed across a variety of measures and relate them. We examined the increase in mean firing rate, the increase in mutual information, and finally how those changes related to variety of modulations observed for neural tuning curves.

Across the population of neurons there was an increase in firing rate and mutual information for saccades towards the RF at saccade onset **(Figure 5)**. In each monkey, we plot the firing rates over time averaged across neurons after normalizing to the peak response. Because Marmoset M was recorded using linear arrays with higher neuron yields, we plot each animal separately to avoid it from dominating a pooled sample. Neither of the monkeys exhibit a strong modulation in rate at the time of stimulus onset but immediately around saccade onset showed an increase in rate for saccades towards the RF **(Figure 5, A-B)**. To quantify the distribution of effects in the population we used an attention index (A.I.) defined as (towards rate – away rate)/(towards rate + away rate). At stimulus onset (not shown) the distribution of AI indices for one monkey was centered near zero reflecting no increase in rate (Signrank test, median −0.3%, AI = −0.002, p = 0.065). In the second monkey (M) there was a modest 3.0% increase (Signrank Test, Marmoset M: AI = 0.015, p = 0.00001). By comparison, as saccade onset both monkeys showed significant increases reflected by a rightward shift in the distribution of AI indices **(Figure 5C)**. At that time Marmoset E showed a 14.2% median increase and Marmoset M showed a 13.8% median increase (Signrank Test, Marmoset E: median AI = 0.0738, p= 1.134e-15; Marmoset M: AI = 0.0615, p = 4.857e-29) and these increases did not differ from each other significantly (Ranksum Test, p = 0.7418). Thus, both animals show a consistent increase in firing rate during the pre-saccadic epoch with similar rapid timing at saccade onset.

**Figure 5.**
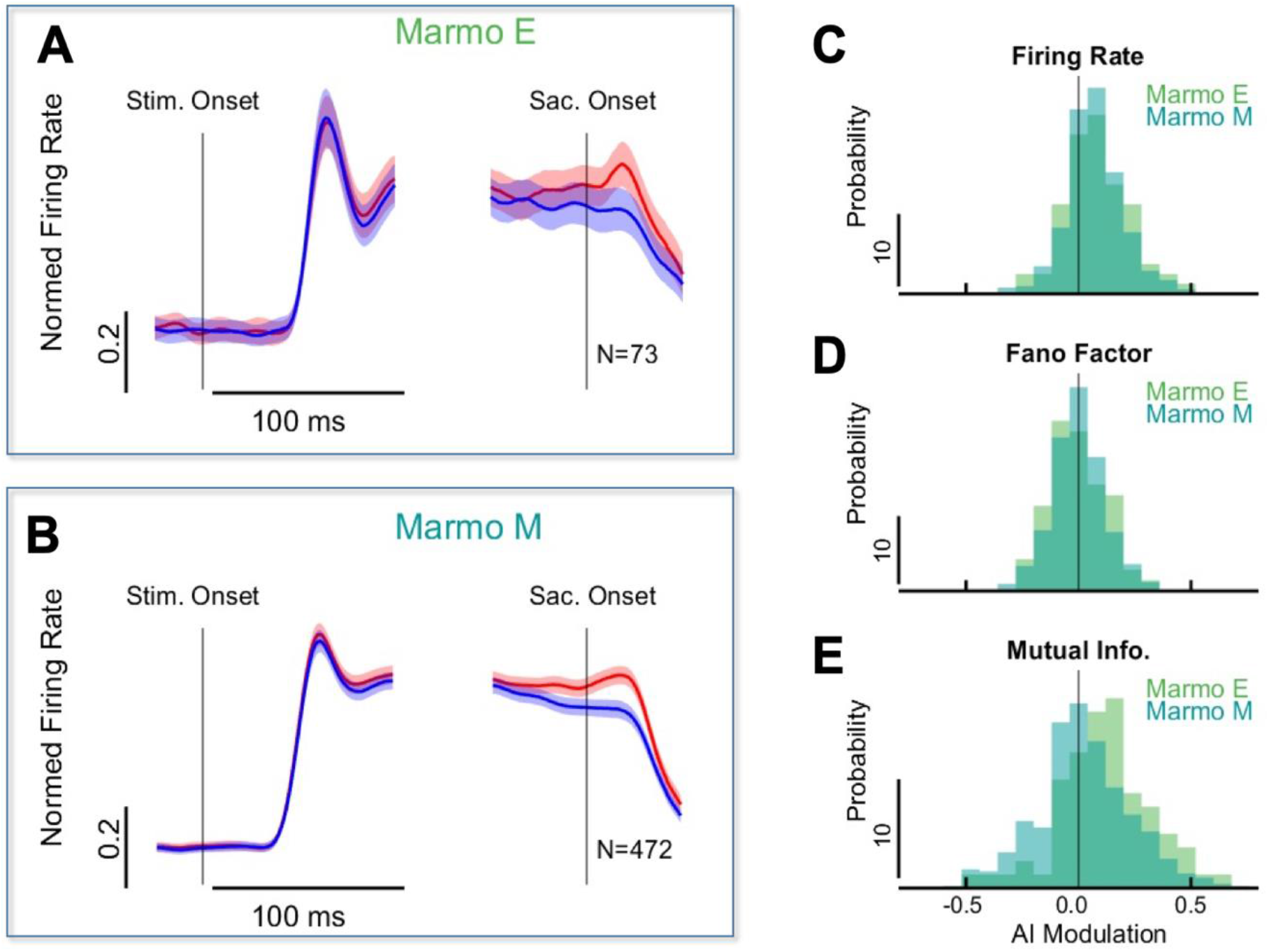
Pre-saccadic attention enhances rate and sensitivity in MT/MTC cells. **(A)** The population mean firing rate for Marmoset E aligned to Stimulus Onset (Left) and Saccade Onset (Right) for the “Towards RF” condition (in red) and the “Away RF” condition (in blue). Error bars are 2 S.E.M across the population. **(B)** The population mean firing rate for Marmoset M (same conventions as in A). **(C)** Histogram of the attention indexes (A.I.s) for firing rate during the saccade onset epoch for Marmoset E (green) and Marmoset M (cyan blue). A rightward shift in the distribution indicates an increase in rate. Histogram of the AI distributions of Fano Factor did not indicate a significant reduction as a leftward shift **(D).** Histograms for mutual information did indicate a rightward shift reflecting increases in neural sensitivity **(E)**.

Increases in firing rate, however, are not necessarily directly correlated to increases in sensitivity as that depends also on whether or not increases are multiplicative or additive to the tuning curve, and how variability in firing is modulated. We therefore examined Fano Factor (FF), a measure of firing variability that is normalized by the firing rate. We found only modest reductions in FF for the two marmoset monkeys **(Figure 5D)** that were not significantly different for Towards vs Away conditions (Signrank Test, Marmoset E: −3.6%, AI = −0.0188, p = 0.595; Marmoset M: −1.2%, AI = −0.0065, p = 0.595). In the second monkey, where we had performed linear array recordings, we examined the shared variability between pairs of neurons, noise correlations, and found a similar modest reduction during the pre-saccadic epoch (Marmoset M: N=21,169 pairs, −4.1%, AI=-0.019, p = 0.012). This result differs from covert attention, where previous studies have found much larger reductions both in Fano Factor and noise correlations (Cohen & Maunsell, 2009; Mitchell et al., 2009).

However, despite the modest reductions in variability, we nonetheless observed consistent increases in the mutual information across both monkeys for the Towards RF as compared to Away RF condition **(Figure 5E)**. Both animals had an individually significant increases, though they did differ in magnitude with Marmoset E having a median increase of 27.33% (Signrank Test, AI= 0.1202, p = 1.5709e-04) and Marmoset M showing a modest increase of 3.9% (Signrank Test, AI = 0.0193, p = 0.0161), a difference between animals that was significant (Ranksum test, p = 0.0015). Thus, while both marmoset monkeys show increases in rate and in sensitivity, which qualitatively are similar to sensitivity enhancements found in macaques, there remains a question about why the magnitude of their effects differ. We thus sought to relate these changes to underlying changes in neural tuning curves, and in doing so also address to what extent either of these animals might show changes consistent with a feature-specific selection in gain.

### Changes in neural tuning enhance sensitivity but not feature-gain

To quantify changes in tuning we fit an adjusted Von Mises function to the motion direction tuning in the saccade towards and away conditions. The Von Mises function fits a single peaked tuning curve to the neural responses as a function of motion direction or orientation (Patterson et al., 2013). It is defined by four parameters: a baseline, an amplitude (gain), a width of the tuning curve, and a preferred direction. To make meaningful comparisons of the tuning between the saccade conditions we limited our analyses to those neurons for which the model fit was better than a minimum R-squared criterion (R^2^ > 0.5, see methods). This included 33 cells from Marmoset E and 284 cells from Marmoset M. Although this sample was biased towards neurons with higher mean firing rates and directional tuning, it allowed more reliable comparison of changes in tuning.

Across the population we found that tuning curves showed increases in either baseline firing or gain but no net change in tuning width **(Figure 6)**. To quantify effects, we computed attention indices (AI = (towards-away)/(towards+away)) for each of the fit parameters. Baseline firing rates increased reflected by a rightward shift in the distribution of attention indices **(Figure 6A)**, with Marmoset M showing an 18.9% median increase that was significant (Signrank Test, AI = 0.0865; p = 1.5010e-07) and Marmoset E approaching significance with an 8.4% median increase (Signrank Test, AI = 0.0404; p = 0.2278). The difference between animals in baseline, however, was not significant (Ranksum test, p = 0.425). The modulation of gain also showed significant increases for both animals **(Figure 6B)** but with a robust increase of 29.5% in Marmoset E (Signrank Test, AI = 0.123; p = 0.0011) and only a modest increase of 5.2% in Marmoset M (Signrank Test, AI = 0.0253; p = 0.0018), which did differ significantly between monkeys (Ranksum test, p= 4.9683e-04). In contrast to baseline and gain, the tuning width exhibited no net increase or decrease in either monkey (Marmoset M: +1.1%, AI = 0.0029; p = 0.9321; Marmoset E: −0.9%, AI = −0.0024; p = 0.5259) which was reflected by attention indices clustered around zero **(Figure 6C)**. Thus, across the two animals increases in baseline and gain contributed to average changes in tuning, while changes in tuning width were not consistent.

**Figure 6.**
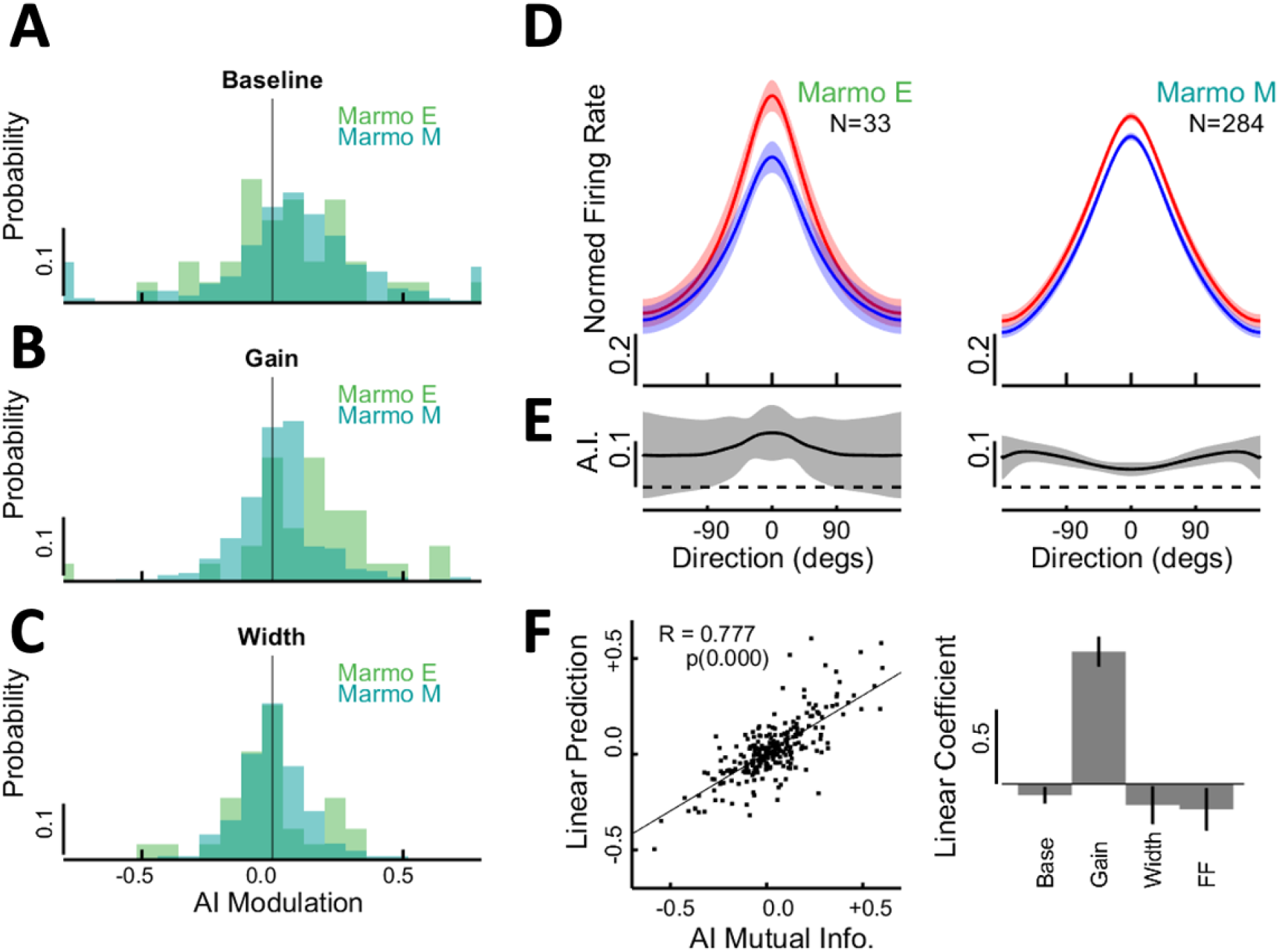
Modulation of neural tuning reflects baseline and gain increases. **(A-C)** The distribution of attention indices (A.I.) are shown for the fit Von Mises parameters of baseline, gain, and tuning width for each animal (Marmoset E, green; Marmoset M, cyan blue). Rightward shifts in the distributions indicate increases in baseline **(A)** and gain **(B)**, with no clear change for tuning width **(C)**. **(D)** The normalized tuning curves broken out by monkey for the “Towards RF” condition (red) versus the “Away RF” condition (blue). Error bars are 2 S.E.M. **(E)** Average gain represented by the A.I. as a function of motion direction for each animal is shown (error bars are 2 S.E.M). **(F)** (Left) Changes in sensitivity as measured by the A.I. in mutual information can be predicted by a linear model with A.I. for baseline, gain, width, and Fano Factor as inputs. Data are pooled across animals. (Right) The prediction coefficients of the linear model for AI Fano factor, AI gain, AI base and AI width (data pooled across both animals). The error bars are 95% confidence intervals on the linear prediction coefficients. Gain provided the dominant prediction of sensitivity.

The average change in tuning between the two animals differed with one monkey showing predominantly increases in gain and the other a stronger baseline increases with modest changes in gain **(Figure 6D)**. We considered to what extent those difference might also reflect changes consistent with feature-specific changes in gain. A spatial modulation would show a consistent increase in gain across all directions regardless of a neuron peak in tuning, while a feature specific modulation for the saccade target would preferentially increase the gain at the peak while suppressing it in the flanks **(Figure 1D)**. Overall, the gain remained uniform across motion directions in both monkeys, more consistent with a spatial rather than feature gain modulation **(Figure 6E)**. Marmoset E shows a weak trend for a larger gain at the preferred direction than at non-preferred direction, but comparing AI values between the preferred and non-preferred directions this difference was not significant (Ranksum, median AI difference = +0.055, p = 0.3669). Meanwhile Marmoset M showed the opposite trend with smaller gain at the preferred direction than the non-preferred direction, a difference that was significant but in the opposite direction predicted by the feature gain model (Ranksum, median AI difference = - 0.049, p = 0.0098). Thus, the changes in tuning largely support a uniform spatial gain, rather than feature gain, which would be more consistent with previous studies of covert attention.

Finally, we examined how well we could predict the observed changes in neural sensitivity based on the underlying changes in tuning. A linear predictor of the attention indices (A.I.) for mutual information (sensitivity) for each neuron was fit using the AI for Fano Factor, baseline, amplitude, and tuning width as input parameters **(Figure 6F, left)**. Although the mutual information is a non-linear measure of sensitivity, it was fit well overall by these variables with an R = 0.764 (Spearman’s rho, p = 1.663e-54). The strength of the linear coefficients in the fit prediction also provides an estimate of their importance for increases in sensitivity **(Figure 6F, right)**. We find that changes in gain predominately drove the changes in sensitivity with the largest prediction coefficient, while baseline and Fano Factor had a weaker and opposite influence. Increases in tuning width also tended to reduce predicted sensitivity but did not reach significance. Overall, changes in sensitivity were primarily driven by increases in gain, and thus the larger increases in sensitivity for Marmoset E can be explained by that animal’s larger increases in gain.

As a final consideration, we performed a control analysis to rule out that behavioral preferences to select the RF location over other away locations could have explained our results. We computed an attention index to represent the location bias for each behavior session as the number of saccades towards the RF minus the number away, normalized by their sum (i.e., an attention index for saccade location preference). We tested for correlation between AI for location preference and each of the tuning parameters (baseline, gain, and tuning width). For each monkey and across all measures there was no significant correlation (Spearman’s rho, p>0.05). For Marmoset M, there was a correlation approaching significance for the baseline parameter (r = 0.083, Spearman’s rho, p = 0.058). We further tested if this correlation might be present in the epoch from 40-100 ms after stimulus onset, where this animal had exhibited a modest increase in mean rate for the Towards RF condition (3.0% increase in rate, p = 0.00001, Figure 5B). Indeed, we did find a significant correlation in that early epoch with location biases in Marmoset M (r = 0.102, Spearman’s rho, p = 0.029). This suggests locations biases influenced early firing in the stimulus epoch for that animal, however, they did account for the larger modulations found in the saccade onset period, which remained similar in magnitude after removing location bias as covariate (Marmoset M: baseline increase, +22.3%, AI = 0.100, p = 1.37e-05). Thus, while location biases may have contributed to early increases in rate for Marmoset M, they did not contribute to the changes reported at saccade onset.

## Discussion

The current findings do not support that pre-saccadic attention automatically engages feature selection for the saccade target, at least not at the level of single units. Recent human psychophysics suggests that pre-saccadic attention differs in important ways from covert attention, one of which involves an automatic feature selection for the target reflected by narrowing in psychophysical sensitivity around its features (Li et al., 2021a; Ohl et al., 2017). At the neural level, studies of feature-based attention suggest there should be an increase in gain at the preferred direction while non-preferred directions are suppressed (Treue & Martinez-Trujillo, 1999; Martinez-Trujillo & Treue, 2004). We measured neural tuning curves for motion direction in areas MT/MTC during the pre-saccadic epoch to test for feature-gain for the saccade target. However, we do not find strong evidence for feature-gain across the total population. While some neurons may exhibit narrowing in their tuning, other show the opposite pattern just as frequently. Overall, the magnitude of gain was largely uniform across motion directions, which is instead consistent with the spatial gain which has been previously observed in studies of covert attention (McAdams & Maunsell, 1999a). While our findings are limited in addressing the read-out of activity at the population level, which may be more closely linked to behavior, we can at least conclude that the patterns of modulation among single neurons are highly similar to that found in covert attention, rather than feature-based attention.

Previous human studies that found feature-specific enhancements employed tasks that additionally require a stimulus discrimination. The foraging task required only that marmosets plan a saccade towards or away from the receptive fields of neurons under study, without any demands to discriminate the target. Thus our task isolates aspects of pre-saccadic attention that are only obligatory to saccade planning. While we found increases in neural firing and sensitivity to confirm the automaticity of pre-saccadic enhancement, we do not find feature-specific enhancements for the saccade target as some previous human studies (Li et al., 2017; Ohl et al., 2017). Those differences may simply reflect that the brain areas we are studying are not involved in those aspects of pre-saccadic attention, or differences between species. However, it is also possible that task differences underlie the lack of feature-specific enhancements. Human tasks involved a discrimination of a grating titled left or right from vertical that flashed briefly within a sequence of 1/f masking noise. Such a task is more likely to engage feature-attention for the reference grating to isolate it from masking noise. While one of these previous studies did show that covert and pre-saccadic attention differed under identical discrimination demands (Li et al., 2016), it is possible that some degree of feature-attention had to be engaged in the task in order to reveal the amplification by pre-saccadic attention. Thus it is possible that feature-specific enhancements are not an automatic feature of pre-saccadic attention.

This is the first study to examine neural mechanisms of attention in the marmoset monkey, a small New World primate. While it is already established that the marmoset shares similar feed-forward circuits as macaques from retina to cortex for visual processing (Troilo et al., 1993; Solomon & Rosa, 2014; Mitchell & Leopold, 2015), as well as frontal-parietal networks involved in eye movement control (Solomon & Rosa, 2014; Gharemani et. al., 2017; Johnston et al., 2018), little is yet know about the mechanisms of selective attention. In the macaque almost everything we know about the neural mechanisms of attention originates from tasks that involve covert attention tasks with long delay periods of sustained central fixation prior to making a judgment or saccade to a peripheral target (Moran & Desimone, 1985; Treue & Maunsell, 1996; Seidemann & Newsome, 1999). Studies of pre-saccadic attention in macaques have also used delayed saccade task with sustained fixation periods before the saccade (Moore & Chang, 2009; Steinmetz & Moore, 2010). Here we found that a saccade foraging paradigm with minimal fixation delays was sufficient to examine neural modulation of pre-saccadic attention in the marmoset monkey. Marmosets completed 400-600 accurate saccade trials in daily sessions enabling us to map neural tuning curves across a range of 16 motion directions. Neurons in marmoset area MT showed neural modulation consistent with that of macaques using more demanding delayed saccade tasks (Moore & Chang, 2009; Steinmetz & Moore, 2010; Steinmetz & Moore, 2014; Merrikhi et al., 2021). Neurons increase their mean firing rate and sensitivity to motion direction in the period immediately prior to saccade onset. This supports that the neural mechanisms of pre-saccadic attention are conserved from Old World to New World primates, and that they generalized to more natural task conditions.

One difference from findings in previous studies of attention in macaques is that we did not find a significant change in firing variability. There was a modest reduction in the Fano Factor in both animals (4.8% and 0.6% respectively), but it did not reach significance. And in the one monkey where we had made array recordings, we also found a modest (−4%) reduction in noise correlations between pairs of neurons, but while significant this reduction was modest relative to the nearly 50% reductions reported in covert attention (Cohen & Maunsell, 2009; Mitchell et. al. 2009). This difference might reflect the emphasis of our experimental design to sample form complete tuning curves, wherein we collected fewer trial repetitions per motion direction. Previous studies have focused on getting larger trial repetitions using fewer stimuli, often only a preferred and/or non-preferred stimulus for each neuron (Mitchell et al., 2007; Li & Basso, 2008; Moore & Chang, 2009; Steinmetz & Moore, 2010). However, these differences could also reflect differences in the behavioral tasks, which would be interesting to explore in future studies. Specifically, while previous paradigms measured variability during extended periods of sustained central fixation (Mitchell et al., 2007; Li & Basso, 2008; Moore & Chang, 2009; Cohen & Maunsell, 2009; Steinmetz & Moore, 2010; Merrikhi et al., 2021) we allowed marmosets to initiate saccades with no delay at stimulus onset. It is known that stimulus onsets can quench noise correlations (Churchland et al., 2010), which could explain the modest reductions in our task where saccades were initiated within 150 to 250ms of stimulus onset. However, it is also worth pointing out that during natural saccade foraging saccades will typically occur in rapid succession every 200-300 ms (Mitchell et al., 2014), so in many ways the timing in our task may better reflect what is relevant during natural vision.

Our study establishes that pre-saccadic attention modulates neural responses in extra-striate cortex of the marmoset monkey, but also reveals a diversity in how neural tuning curves are modulated. Recent evidence from covert attention studies demonstrates that depending on the read-out, information can be reshaped in the read-out to enhance specific types of information, even while there may be no net improvement the total population (Ruff & Cohen, 2019). Distinctions in how different parts of the population encode information are supported by laminar distinctions found in other studies of covert attention (Buffalo et al., 2011; Nandy et al., 2017; Pettine et al., 2019). And a recent study of pre-saccadic attention found that stimulus orientation was better encoded by superficial layers neurons whereas movement direction was better encoded by deep layer neurons (Pettine et al., 2019). We find substantial variety in the way neural tuning curves are modulated, and thus depending on how information is read-out, it remains possible that changes in tuning with pre-saccadic attention could support a feature-selection of the target in the read-out of population activity. However, at the level of individual units, pre-saccadic modulation appears consistent with changes reported in covert attention.

## Acknowledgments

We would like to thank Dina Graf and members of the Mitchell lab for help with marmoset care and handling. We thank Martin Rolfs and Lisa Kroell for comments on an earlier draft of this manuscript. This work was supported by NIH grants R01 EY030998 (JFM and SC), R00 EY032179 (JY), and T32 EY007125 (SC).

## Methods

### Subjects and Surgery

All experimental protocols were approved by the University of Rochester Institutional Animal Care and Use Committee and were conducted in compliance with the National Institutes of Health guidelines for animal research. Two adult common marmosets (Callithrix jacchus), Marmoset E (female) and Marmoset M (male), were used for neurophysiology recoding experiments to measure changes in neuronal tuning during presaccadic attention. Subjects were single housed at the University of Rochester with a circadian cycle of 12-hour light and 12-hour dark. Subject M was briefly food scheduled with full access to water during early training but had no restrictions by the time neurophysiological data was collected. Subject E was never food scheduled and always had full access to food and water.

Both subjects were surgically implanted with head caps to stabilize them for head-fixed eye tracking and neural recordings. Two months prior to surgery, subjects were trained to sit in a small primate chair following methods previously described (Lu et al. 2001; Remington et al. 2012; Osmanski et al. 2013; Nummela et al. 2017). Then subjects underwent surgery under sterile conditions to implant an acrylic head cap with titanium posts to stabilize the head using methods described in detail previously (Nummela et al., 2017). During the implant surgery, recording chambers were placed over visual areas MT and V1 based on stereotaxic coordinates (Paxinos et al., 2012). Recording chambers consisted of custom 3D prints (Proto Labs) and adhered to the skull using C&B-Metabond (Parkell, Inc.). The skull inside the recording chambers was also covered by a thin layer of C&B-Metabond. After initial head-implant and chamber placement, marmosets were trained to acclimate to head-restraint while sitting comfortably in a custom designed primate chair. Over several months they were trained to perform several basic tasks including central fixation (Mitchell et. al., 2014) and a saccade task towards a peripherally detected Gabor grating which we used to measure their visual acuity (Nummela et. al., 2017).

After preliminary training a second surgery was performed to create a craniotomy (2-3 mm in diameter) in the recording chamber over area MT. Craniotomies were sealed with a thick layer of silastic gel (Kwik-Sil; World Precision Instruments) to protect the brain from infection and reduce granulation growth (Spitler & Gothard, 2008). If any bleeding occurred or the Silastic seal leaked clear fluids in the days following surgery, then the chamber was cleaned with sterile saline and a new Silastic layer applied. Typically, the chamber stabilized and remained dry with a tight seal after a few days to a week. At that time, we performed dural scrape to remove any excess tissue over the dura and applied a thin layer (<1mm) of Silastic which was thin enough to enable passage of electrodes for recordings. The silastic remained in place for the duration of the study. When applied correctly, the silastic has been observed to limit the growth of granulation tissue on the dura and prevent infection (Spitler & Gothard, 2008), and can also be recorded through with tungsten electrodes (Miller et al., 2015, MacDougall et al., 2016). Additionally, it was possible to record through Silastic using linear array silicon probes (NeuroNexus, Inc.) when the dura was thin.

### Electrophysiology

Over the duration of the study our recordings improved from using single channel tungsten electrodes into using multi-channel linear silicon arrays. The first monkey was recorded entirely with tungsten single electrodes (47 sessions), with fewer tungsten electrode recordings in the second monkey (31 sessions). The bulk of data from the second monkey originated from linear arrays that provide much higher cell counts per session (21 sessions). Due to animal health issues, we were unable to employ linear arrays in the first monkey which passed away during the pandemic. The current findings focus on behavior and the single unit neural effects on tuning curves during presaccadic attention, which can be addressed well with either recording method and including both animals.

In single tungsten electrode recordings, we sampled across recording sites in a 1mm spaced grid with electrodes advanced through metal guide tubes placed in the grid. We inserted tungsten 2.5- to 5-MΩ electrodes (1-3 FHC) that were mounted onto a lightweight screw micro-drive (Crist Instrument, 3-NRMD drive) positioned over the spacing grid. Electrodes were passed through a metal guide tube that touched but did not penetrate the Silastic layer covering the brain. Tungsten electrodes reliably penetrated the thin silastic layer and intact dura to enter the brain. A stainless-steel reference wire was implanted under the skull in a 1mm craniotomy.

Later recording sessions using multi-channel linear silicon arrays (NeuroNexus, Inc.) used a custom-built X-Y stage for sub-millimeter targeting of recording sites. The X-Y stage mounted onto the recording chamber and carried a light-weight screw micro-drive (Crist Instrument, 3-NRMD) that could deliver linear arrays mounted to a steel tube (28 gauge) into the brain. Design for the 3D printed parts used in the X-Y stage and recording chamber are online at https://marmolab.bcs.rochester.edu/resources.html).

All neurophysiology data was amplified and digitized at 30kHz with Intan headstages (Intan) using the open-ephys GUI (https://github.com/open-ephs/plugin-GUI). The wideband signal was highpass filtered by the headstage at 0.1 Hz. We corrected for the phase shifts from this filtering (Jun et al., 2017). For linear arrays, the resulting traces were also preprocessed by common-average referencing.

### Tungsten spike sorting and cluster isolation

Single-unit and multi-unit clusters from tungsten electrodes were identified using custom MATLAB software. First, the raw signal was band passed filtered from 800 Hz to 6000 Hz with a 6th-order Butterworth filter. To reduce movement artifacts in recordings (i.e., licking, or other movements), we choose a narrower filter for initial spike detection to threshold spike events (1500 Hz to 4500 Hz), and then used the wider band pass (800-6000hz) to classify single units based on clustering of their threshold trigged spike waveforms in a principal components analysis (PCA) space that included the first two principal components and time as variables. Clusters identified in PCA space were compared to a noise floor, based on random sampling of threshold events, and clusters that could not be fully separated from other clusters or the noise floor, or that exhibited more than 1% of inter-spike interval violations under 1 ms, were counted as multi-unit activity.

### Laminar electrophysiology and spike sorting

In later recordings, we were able to use multisite silicon electrode arrays that provided much higher cell counts. These arrays included 1-2 shanks and each shank consisted of 32 channels with 35 microns spacing between contacts. All arrays were 50 microns thick and had sharpened tips. Arrays that included two shanks spaced 200 microns apart were from NeuroNexus (http://www.neuronexus.com). Although the arrays could penetrate dura when it was thinned, we found that dimpling of the tissue could still suppress neural activity during insertion and that the best recording quality was achieved by applying a small 1-2mm horizontal slit in the dura during a dural scape and sealing it under Silastic to prevent infection. For the best recording quality, it was further useful to electrode-plate the silicon electrode arrays with PEDOT, a method that has been shown to increase signal/noise ratios (Ludwig et al., 2006 & Ludwig et al., 2011). Last, the yield of neurons recorded was generally improved by inserting the array electrodes into cortex slowly. We first lowered the silicon shank quickly until we observed units at the array tip, and then retracted one turn of the micro-drive (250 micron) slowly. Then we lowered the arrays advancing approximately 4-6 turns (1 to 1.5mm) over a 20-30 minute duration until neurons were evenly distributed across the length of the shank. We then slowly retracted the array 1-2 turns (0.25-0.5 mm) to reduce pressure on the tissue during recordings. During this final retraction, neurons did not typically shift vertical locations on the array, suggesting that it primarily acted to reduce pressure on the tissue and that otherwise the arrays would have continued advancing slowly during the recording as the tissue relaxed. After retracting, we waited 20 minutes before starting the main behavioral task and recordings.

We spike sorted array data after initial filtering from the INTAN system (as described earlier) using Kilosort2. Outputs from the spike sorting algorithms were manually labeled using the ‘phy’ GUI (https://github/kwikteam/phy). Units with tiny or physiological implausible waveforms were classified as noise and excluded. Kilosort can identify multi-unit clusters that are not physiologically possible, spanning any channels with unrealistic waveforms. Therefore, to be conservative we only included units from Kilosort that had clear clusters in PCA space, less than 1% inter-spike interval violations, and bi-phasic spike waveforms localized to adjacent channels on the linear array.

### Stimulus presentation and timing

Stimuli were generated using the Psychophysics toolbox (Brainard, 1997; Kleiner et al., 2007; Pelli, 1997) in MATLAB 2015b (MathWorks, Natick, MA) on a PC computer (Intel i7 CPU, Windows 7, 8 GB RAM, GeForce Ti graphics card). They were presented on a gamma-corrected display (BenQ X2411z LED monitor, resolution: 1,920 x 1,080 p, refresh rate: 100Hz, gamma correction: 2.2) that had a dynamic luminance range from 0.5 to 230 cd/m^2^ at a distance of 57 cm in a dark room and viewed under head-restraint in custom designed primate chair as described previously (Nummela et. al., 2017). Brightness on the display was set to 100 and contrast to 50, and additional visual features of the monitor, such as blur reduction and low blue light, were turned off. Gamma corrections were verified with measurement by a photometer. Task events and neural responses are recorded using a Datapixx I/O box (Vpixx technologies) for temporal registration. Matlab code is available online (https://github.com/jcbyts/MarmoV5).

Random dot motion fields were used as targets for saccade foraging and also provided a stimulus to validate the motion selective responses on individual neurons as an inclusion criteria. Each aperture contained a field of black dots (each dot 0.15 dva diameter with a density of 2.54 dots per visual degree squared) which moved at 15 degrees/sec in one direction (100% coherent). The dots had limited lifetimes of 50 ms with asynchronous updating to new locations. The radius of the dot field was equal to half the eccentricity of where it was located from the center of the screen and appeared over a gray background (115 cd/m^2^). The contrast of dots was decreased from black at the center of field (0.5 cd/m^2^) to zero on the aperture edges according to a Gaussian envelope with a sigma equal to a 1/6^th^ of the aperture’s diameter.

### Eye Tracking

Eye position was acquired at 220 Hz using an Arrington Eye Tracker and Viewpoint software (Arrington Research) or at 1000 Hz using an Eyelink 1000 Plus eye tracker (SR research). Eye position was collected from infrared light reflected off of a dichroic mirror (part #64-472, Edmunds Optics). Each subject’s vision was corrected using spherical concave lenses (Optimark Perimeter Lens Set) that were centered 4–5 mm in front of the face as described previously (Nummela et al. 2017). The lens of −2.5 diopters was used for marmoset M and of - 2.0 diopters for marmoset E. Eye position was calibrated at the start of each behavioral session using a Gabor windowed face detection task described previously (Mitchell et al., 2014, Nummela et al. 2017).

Eye-position data was collected during the entire recording session. Raw horizontal and vertical eye position signals were smoothed offline with a median filter (5 samples, 5 ms) and convolved with a Gaussian kernel (5 ms half width, out to 3 SD, −15 to 15 ms) to minimize high-frequency noise. For off-line detection of saccadic eye movements, we used an automatic procedure that detected deviations in 2-D eye velocity space (Engbert & Mergenthaler, 2006; Kwon et al., 2019). We computed horizontal and vertical eye velocity by taking the difference of the smooth eye-position and then marked saccades by where the 2D velocity exceeded the median velocity by 10 SD for at least 15 ms (Engbert & Mergenthaler, 2006; Kwon et al., 2019) and merged any two saccadic events into a single saccade if they were separated by less than 5 ms. Saccade onset and offset were determined by the first and last time the 2-D velocity crossed the median velocity threshold. Epochs with eye blinks (based on pupil size) are removed from analysis.

### Receptive field mapping of MT/MTC during free viewing task

Spatial receptive fields were estimated from the responses to a wide field stimulus consisting of large moving white dots **(Figure 2A)**. In brief, marmosets freely viewed a full-field display that consisted of white dots (230 cd/m^2^, 1 dva diameter) that appeared against a gray background (115 cd/m^2^) spanning ±20 dva on the horizontal and 15 ±dva on the vertical of the display. Each dot moved at 15 degrees per second for 50 ms before being replotted to a new location in the full field display. Dot motion was selected at random from 1 of 16 motion directions sampled around the circle. In each task trial, the screen would contain a fixed number of dots (sampled from 4,8,16, or 32) that would be viewed for 10 seconds. To encourage foraging near the center of the screen a Gabor target (20% Michelson contrast, 1 dva diameter, 1 cycle/degree, with random orientation) appeared within 5 degrees of the center superimposed with the dots flashing dots. If the marmoset’s eye position acquired the Gabor target within a 2-degree diameter a juice reward was delivered, and the Gabor target was replotted to a new location.

Offline we corrected for eye position to represent the flashed stimuli in a grid of retinal coordinate locations and correlated the stimulus history with spike counts to estimate the receptive field **(Figure 2B)**. The full methods for estimating receptive fields have been described previously (Yates et. al., 2021) and analysis code is available online (https://github.com/VisNeuroLab/yates-beyond-fixation). In brief, firing rate was computed as a function of the x,y retinal-based grid location (2×2 degree bins) where each bin contained a flashed dot or did not. To reduce correlation in the stimulus history of the moving dots, we only represented dots on the first video frame from their 50 ms lifetime, and the spatial position was registered by their location at the middle of the lifetime. The firing rate was computed for the onset of dots across the grid at different lag times in 10ms spike counting bins **(Figure 2B left)**. Those locations exhibiting firing responses significantly above the pre-stimulus baseline firing (from −100 to 0 ms lag, p < 0.001) were labeled as significant to mark the receptive field (RF). A smoothed 2D contour was computed to circumscribe the peak of the RF at its half-maximum height relative to baseline for the peak temporal lag **(Figure 2B, top right)**. Then the direction selective evoked responses were computed from the flash events inside the defined RF contour as a function of their motion direction, with error bars indicating two standard errors of the mean at each direction **(Figure 2B bottom)**.

The tuning for motion in full-field mapping and also later for dot-field stimuli in foraging were fit using a modified von Mises function. The tuning curve for firing rate *R* was defined as a piecewise continuous function based on a bandwidth parameter, *K*, as

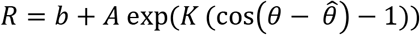

when *K* > 0 and otherwise as

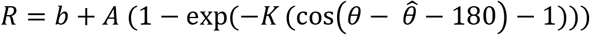

where *b* is the baseline firing rate, *A* is the amplitude, *K* is the bandwidth and 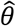 is the preferred direction. Von Mises functions have been used previously to describe motion tuning in area V1 (Patterson et. al., 2013). We adapted that function to allow for curves with wider than cosine tuning. Cosine tuning in the function occurs as *K* approaches 0. We allowed the curve to continue to be defined for negative values of *K* adopting an inverted von Mises with the opposite direction preference (180 degrees opposite) such that the peak location remained at the same preference but was wider than cosine tuning. The modified curve was fit by maximizing the likelihood of the spike counts observed for each motion direction bin assuming Poisson firing statistics (Truccolo et al., 2005). Error bars were generated by a 10-fold Jacknife procedure, and tuning width was estimated from the half-width of the curve in degrees.

### Neural inclusion criteria

Areas MT and adjacent MTC were identified in targeted recordings based upon their direction selective responses and retinotopy (Rosa & Elston, 1998). While we targeted neurons in area MT, a small number of motion selective neurons from adjacent MTC (also called V4t in lower hemi-field and MST-lateral in upper hemi-field for the macaque) may have been included that lied near the vertical meridian. MTC has been reported to contain neurons with similar sized RFs and a significant portion of those cells also have motion selective responses (Rosa & Elston, 1998). We used the response to random dot motion patches placed inside the RF during the foraging task (irrespective of saccade condition) to evaluate if neurons had significant visual and motion selective responses. We only included neurons if they had a significant visual response defined by an increase in spike counts from 50-100 ms following onset of a dot motion stimulus (averaged across sampled directions) as compared to a baseline from −100 to 0 ms before onset (Signrank test, p<0.05). We also required that neurons had a significant direction selective index (D.S.I) indicative of motion tuning. The DSI was computed from spike counts from 50 to 150 ms after the stimulus onset as where

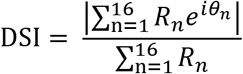

R_n_ represents the mean spike count in response to a motion direction. To compute confidence intervals on the DSI we used a 10-fold Jacknife procedure and units were included if the lower bound on the 95% confidence interval was above a DSI of 0.05.

### Saccade foraging task

To study presaccadic attentional modulations, marmoset monkeys performed a saccade-foraging task in which a saccade was performed to one of three equally eccentric peripheral motion apertures **(Figure 1A)**. In each session, one of the apertures was positioned to fall near the center of the receptive field of a neuron or the set of neurons under study, while the other two were placed 120 degrees apart from it at the same eccentricity. The task trial began with fixation of a small spot (0.3 degree radius, 0.5 cd/m^2^ center, 230 cd/m^2^ surround) within a 1.5 degree radius window for a delay uniformly distributed between 0.1 to 0.3 s, presented on a gray background (115 cd/ m2). After the fixation period, the fixation point was offset and three dot motion apertures (as described earlier) appeared in the periphery. The monkey was given up to 1.5 s to make a saccade out of the fixation window to one of the apertures with the final eye position remaining within the target window for 0.25 sec to confirm the saccade endpoint. We rewarded saccades to any location as long it differed from the previous trial to encourage foraging. A correct choice was rewarded with 10–20 micro liters of liquid reward and the appearance of a marmoset face at the aperture location for 1 s, providing positive feedback. The juice reward consisted of marshmallows blended with water that were prepared fresh for each daily session. An incorrect choice back to a location sampled in the previous trial resulted in a black Gaussian spot filling the chosen aperture, as feedback for choosing the wrong location. The next trial proceeded at an interval of 1-2 seconds depending on juice rewards.

### Trial inclusion criteria

We limited analyses to trials in which saccades to foraging targets were executed in a single step without preceding movements and after a minimum latency from stimulus onset. First, we detected any micro-saccades of amplitude greater than 0.5 visual degrees that occurred during the fixation period and excluded those trials. To ensure that the animal had time to see the stimulus before initiating their saccade, we excluded trials where the reaction time (saccade onset from stimulus onset) was smaller than 0.12 sec. We also excluded trials where the animal made two smaller saccades that stepped to the aperture and trials in which the saccade end-point fell short of the aperture center (> 50% of the target eccentricity). Last, we require that saccades fell within a window that had a radius 50% of the eccentricity of the center of the aperture in order to be counted within that aperture. When all criteria were applied, this resulted in excluding 21.4% and 13.1% of trials across sessions for Marmoset E and Marmoset M respectively. We included a small percentage (10%) of catch trials that required the monkey to maintain central fixation. In these trials, the fixation period was extended to 0.5 seconds without any target apertures ever appearing. For holding fixation, the monkey was rewarded with both juice reward and a marmoset face that appeared at fixation.

### Temporal separation of the stimulus and saccade onset epochs

We examined the neural firing response time-locked relative to the moment of stimulus and saccade onset. Because there is no extended delay period between stimulus and saccade onset under natural foraging tasks, as in a delayed saccade task, these two intervals will be partly overlapping depending on the saccadic reaction time and it is important to choose analysis intervals that minimize their overlap. To examine the temporal response of neurons the peri-stimulus time histograms (PSTHs) for the “Toward” and “Away” conditions were smoothed using a Gaussian temporal kernel (*σ* = 5 ms). The stimulus locked onset response revealed a visual latency around 40ms with a transient peak that rose and fell by 70-80 ms into a sustained response **(Figure 4A,B, left)**. We defined a stimulus response epoch to be between 40 to 90 ms after stimulus onset to capture the early peak. The response when instead time-locked to saccade onset reveals a rise in firing rate continuing up to 20 ms after the saccade followed by suppression **(Figure 4A,B, right)**. We defined the presaccadic window to be between −30 to +30 ms from saccade onset. While previous studies using delayed saccade paradigms have used windows that begin from −100 ms, in our paradigm that interval would include trials with shorter saccadic latencies that include the stimulus evoked peak response. It might be difficult to identify the stimulus evoked response, as it would be convolved with the variability in reaction times, but it would still influence firing rate in those epochs preceding saccade onset. We thus restricted analyses to reaction times later than 120 ms and set our pre-saccadic window to be no earlier than −30 ms from saccade onset such that no part of the stimulus evoked transient, which resolved to a sustained level by 90ms, would be mixed with the saccade onset response. For the end of the saccade onset epoch, we choose +30ms because the visual latency of neurons was no earlier than 40 ms, and thus responses out to that period would yet reflect visual motion induced by the saccade itself.

Although temporal intervals were selected to minimize overlap between stimulus and saccade epochs, we must also control for any differences in saccade reaction times between “Towards” and “Away” conditions when comparing modulation in their firing rates or tuning curves. For example, shorter reaction times in one condition could lead to differences in the adaptation state of the neuron, which if not corrected for, could produce differences in firing at the time of saccade onset. To control for those differences, we created resampled trial distributions for the “Towards” and “Away” conditions that matched the saccadic reaction times between conditions. For each session the distribution of reaction times was computed in 10 ms bins and stepping through each bin we randomly sampled an equal number of trials from the condition with more trials in order to match the number to the condition with fewer trials. An example of a single matched distribution with trials sorted in ascending order by reaction time is illustrated in the raster plot of **Figure 3A**. Ordering the raster plots based on reaction times helps visualize possible influences from the stimulus epoch on the saccade onset epoch. There are no clear signs of the stimulus evoked response contaminating the saccade onset epoch, nor of differences in adaptation between conditions. All comparisons between “Towards” and “Away” conditions applied this correction to be conservative, although all effects reported remained consistent without it throughout the results.

### Firing rate and variability analyses

The mean and variability of firing rate were assessed in the stimulus and saccade onset intervals. The mean firing rate was computed from all trials for either the “Towards” and “Away” conditions, and thus for each of those conditions was sampled randomly over the 16 motion directions used in the task. To measure variability, we computed the Fano Factor (FF), which provides a measure of variance in spike counts across trials that is normalized by rate. For a Poisson process, the spike count variability scales in proportion to the mean count, giving unity FF. However, non-linearities such as the spike refractory period, burst firing, or super-Poisson fluctuations in rate produce dependencies on how the FF scales with rate that deviate from linearity. Thus, it is necessary to match firing rates between the “Towards” and “Away” conditions before comparing them in order to disassociate changes in FF from changes in mean rate (Mitchell et. al., 2007; Churchland et. al., 2010). To match firing rates, we first computed the mean spike counts for each of the 16 motion directions in each of the two saccade conditions (16 points of mean versus variance for each condition). We performed a search in random order for each of the 16 points in the “Towards” condition to find the point with mean rate in the “Away” condition most closely matching its mean rate, without reference to whether motion direction matched, and accepted those as a pair if the two rates matched within 5%. Repeating this procedure without replacement across all points identified the region of overlap between the two distributions that was constrained to have nearly matching rate (< 5%). The FF was then computed by dividing the spike count variance by the rate at each point and then averaging those ratios for each saccade condition. In one monkey where we performed recordings using linear arrays it was also possible to isolate many simultaneously recorded pairs of neurons in order to estimate the noise correlations. Spike counts were computed during the pre-saccadic epoch from −30 to 30 ms at saccade onset and correlations were computed within trial sets where the same motion direction was in the receptive field, and then pooled across motion directions.

### Mutual Information analysis

In the saccade onset interval we examined if there was an increase in neural sensitivity for motion direction. We measured sensitivity from the distributions of firing rates across the 16 motion directions by computing the mutual information. The mutual information quantifies how much information one variable (such as firing rate) provides about another variable (such as motion direction) measured in bits. For our purposes it provides an estimate of sensitivity, much as classic measures like the area under the curve (AUC) in receiver-operator curve (ROC) does for the case of discriminating two sensory conditions, but it is readily generalized to more than two stimulus conditions and thus is highly suitable for use with tuning curves (Hatsopoulos et al., 1998). We calculated a mutual information (MI) for both the “Towards” and “Away” conditions using the equation:

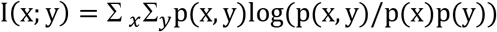

where x is the motion direction stimulus shown and y is the measured firing rate. Computing the mutual information requires that we estimate the probability distributions (p(x), p(y), and p(y|x), using histograms and binning of firing rate data per stimulus direction. For the stimulus variable x we binned based on the motion direction, where we had sampled from 16 possible directions. However, to further enforce a smoothing constraint on our data and represent that adjacent motion directions are related, we further pooled the firing rate data from the two adjacent motion direction bins arranged on the circle, thus giving on average three times as much firing rate data per motion direction bin and forcing a smoothing constraint on the data. The binning of the firing rate, y, was determined based on a “goodbins” function described in Scott (1979), applied to the total firing rate distribution across all motion directions to estimate the marginal distribution p(y). Then the histogram of firing rates conditioned on each stimulus direction, p(y|x), was computed using those bins that were established to describe p(y).

### Comparing motion tuning between towards and away conditions

To examine neural tuning changes across the “Towards” and “Away” conditions, we fit the motion direction tuning of individual MT units with a modified von Mises function as described earlier in methods. The von Mises curves provide parameters for amplitude, baseline, and width for each tuning curve in the two conditions. We constrained fits to share the same preferred direction between saccade conditions. Only units where the net quality of the curve fits for the two conditions (pooled over both curves) exceeding an R-squared of 0.5 were included. This significantly reduced our neural population included for von Mises based analyses: (marmoset E had 33 units and marmoset M had 287 units).

To quantify the modulation of parameters between the “Towards” and “Away” conditions, we calculated an Attention Index (AI) for each of the parameters considered as:

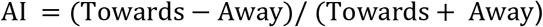

where AI ranges from −1 to 1, with zero indicating no change. The AI metric is useful for emphasizing percentage changes in a variable. All statistical tests on AI indices were non-parametric (Signrank or Ranksum tests, p < 0.05).

### Linear prediction model

To evaluate which modulations in tuning curves contributed to increases in neural sensitivity we fit a linear model. The input parameters to the linear regression included the attention index (AI) for the Von Mises fit parameters of baseline, gain, and curve half-width, as well as the AI for the rate matched Fano Factor. We also include a constant term, giving 5 input parameters. The output aimed to predict the AI for mutual information of each unit. Linear fit coefficients (and error bars) were fit using the ‘regress’ function in Matlab (version 2018). Only units which had reliable Von Mises fits for both the “Towards” and “Away” conditions (R^2^= 0.5) were included.

